# Toxicity and bioaccumulation of Cadmium, Copper and Zinc in a direct comparison at equitoxic concentrations in common carp (*Cyprinus carpio*) juveniles

**DOI:** 10.1101/717363

**Authors:** Vyshal Delahaut, Božidar Rašković, Marta Satorres Salvado, Lieven Bervoets, Ronny Blust, Gudrun De Boeck

**Affiliations:** University of Antwerp - Faculty of Sciences, Department of Biology, Groenenborgerlaan 171, 2020 Antwerp, Belgium; University of Belgrade - Faculty of Agriculture, Institute of Animal Science Nemanjina 6, Zemun, 11080 Belgrade, Serbia

## Abstract

The individual toxicity and bioaccumulation of cadmium (Cd), copper (Cu) and zinc (Zn) towards common carp juveniles was evaluated in a direct comparison in two experimental setups. First, the fish were exposed for 10 days to different metal concentrations. Accumulated metals were quantified and showed a positive dose dependent uptake for cadmium and copper, but not for zinc. Toxicity was in the order Cd>Cu>Zn with 96h LC_50_ values (concentration where 50% of the animals dies within 96h) for Cd at 0.20±0.16 μM, Cu at 0.77±0.03 μM, and Zn at 29.89±9.03 μM respectively, and incipient lethal levels (concentration where 50% of the animals survives indefinitely) at 0.16 μM, 0.77 μM and 28.33 μM respectively. Subsequently, a subacute exposure experiment was conducted, where carp juveniles were exposed to 2 equitoxic concentrations (10% and 50% of LC_50_ 96 h) of the three metals. The gill metal content was quantified after 1, 3 and 7 days, and was correlated to electrolyte levels and structural damage of the gill tissue and associated pathological effects. Again a significant dose-dependent increase in gill cadmium and copper, but not in zinc, was observed during the 7-day exposure. Copper clearly affected the sodium levels in the gill tissue, while zinc and cadmium did not significantly alter any of the gill electrolytes. The overall histopathological effects (e.g. hyperemia and hypertrophy) of the metal exposures were mild for most of the alterations, and no metal specific pattern was elucidated for the tested metals except oedema of the primary epithelium which typically occurred in both levels of Zn exposure.

## Introduction

Anthropogenic input of metals remains one of the major threats towards aquatic animals and entire natural ecosystems. These contaminants may originate from a wide range of sources, i.e. industrial activities, traffic, households and agriculture (Wood et al. 2012a; Wood et al. 2012b). Amongst others, cadmium (Cd), copper (Cu) and zinc (Zn) are three metals which are often encountered in surface waters at levels exceeding the environmental quality standards set by the European Union (Commission 2008). In Belgium, total concentrations of cadmium, copper and zinc measured by the Flemish environmental agency (VMM) in 2014, ranged respectively from 4.45e^-4^-0.03 μM (0.05-3.37μg/L), 0.02-0.54 μM (1.27-34.32 μg/L) and 0.12-5.05 μM (7.84-330.17μg/L) (VMM 2014).

Fish living in metal polluted environments might either be exposed to the metals through the food chain, or via direct uptake from the contaminated water. In the latter case, the gills are the first organs to suffer from this kind of pollution and will show the first clinical signs induced by waterborne metal exposure (Guo et al. 2018). Copper and zinc are essential nutrients for fish, and therefore dietary or waterborne intake of these elements is necessary to sustain basic metabolic processes, in contrast to xenobiotic metals such as cadmium (Wood et al. 2012a). However, elevated concentrations of copper or zinc will also lead to adverse effects on a wide range of crucial pathways.

Copper uptake is facilitated via two distinctive mechanisms: a transmembrane protein (copper transporter 1), insensitive to external copper concentrations, and the apical Na^+^-uptake pathways located at branchial epithelial cells, sensitive to external concentration of copper (Grosell and Wood 2002; Mackenzie et al. 2004). In the latter case, intracellular sodium levels can decrease as a direct consequence of competition at the uptake site (Niyogi et al. 2015). In addition, once they enter epithelial cells, copper ions are showing an ability to inhibit the activity of the membrane bound Na^+^/K^+^-ATPase (Li, et al. 1996; De Boeck et al. 2001). Further, copper is known to induce oxidative stress, olfactory impairment, increased plasma ammonia and disturbed acid-base balance (De Boeck et al. 2007; Green et al. 2010, Eyckmans et al., 2011). Acute toxic concentrations of zinc are generally higher than those of copper, yet deleterious effects have been described repeatedly (Hattink et al. 2006; Malekpouri and Asghar 2011). Unlike copper, zinc uptake is facilitated via a common zinc-calcium transport carrier, located in the mitochondria rich cells of the branchial apparatus (De Schamphelaere and Janssen 2004; Mebane et al. 2012). An impaired branchial Ca^2+^-influx and hypocalcemia as an indirect consequence is therefore to be expected (Spry and Wood 1985; Niyogi and Wood 2003).

Prolonged elevated zinc uptake will eventually lead to critically high accumulation in tissues where it can generate damaging reactive oxygen species (Loro et al. 2012). Cadmium, on the other hand, is not essential for fishes’ metabolic processes, and is potentially dangerous at lower concentrations compared to the essential metalloids. As it is the case for zinc, cadmium utilizes the Ca^2+^-channel to enter the gills and can affect calcium homeostasis as well (Suzuki et al. 2004). Once accumulated in other tissues, it will also induce oxidative stress through the generation of reactive oxygen species at relatively low concentrations (Almeida et al. 2002; Reynders et al. 2006; Casanova et al. 2013).

Exposure of freshwater fish to elevated metal levels in the aquatic environment, and the subsequent disturbance in iono- and osmoregulatory processes, typically results in morphological changes of their branchial apparatus (Fonseca et al. 2016; Yancheva et al. 2016). Due to the high plasticity of gill tissue, abovementioned physiological mechanisms can induce tissue damage and/or remodeling. Secondary lamellae of gills serve as a primary site for gas exchange and ion transport in fish, and morphological adaptations either facilitate oxygen uptake or serve as a mechanism for increasing the blood-water barrier (Nilsson et al. 2012). If the presence of the irritant is persistent, different histopathological alterations can occur, which could reduce the respiratory surface and impair respiration and physiology of the gills (Phuong et al. 2017). These histopathological alterations could be quantified to assess adverse effects of specific xenobiotics on the gill tissue, thus giving an insight to the general fish health. Such a methodology is used in a vast number of studies of metals on gill histology, i.e. laboratory exposures (Vuorinen et al. 2003; Suiçmez et al. 2006; Mishra and Mohanty 2008); semi-field set-ups (Schwaiger et al. 1997; Nimick et al. 2007); and field studies (Triebskorn et al. 2008; Fonseca et al. 2017; Kostić et al. 2017). A specific pollutant will not necessarily induce pollutant-specific histopathological alterations in the gill tissue, but a quantification of the extent and intensity of overall changes enables a comparison between experimental groups in various trials (Bernet et al. 1999; Nunes, Antunes, et al. 2015; Nunes, Campos, et al. 2015). This method is utilized in the present study, in order to compare the effects of the three metals on the morphology of the branchial apparatus of common carp.

Exposure concentrations may either be chosen based on the outcomes of environmental studies or come from standard toxicity tests. In the latter case, the toxicity of a substance can be quantified by determining the concentration at which 50% of the exposed individuals die over a time span of 96 hours (96h LC_50_). However, even under controlled conditions the outcomes of these kind of studies are likely to be influenced by confounding factors, such as age of the animals, environmental conditions, and the water chemistry (Stouthart et al. 1996; Das and Das 2005; Ebrahimpour et al. 2010). A query in the EPA ecotox database (EPA 2016) for 96h LC_50_ values of copper, cadmium and zinc toxicity for common carp, resulted in a wide range of concentrations for each of the metals tested: 96 h LC_50_ for Cu between 0.02 and 542 μM (Rehwoldt et al. 1971; Deshmukh and Marathe 1980; Verma et al. 1981; Alam and Maughan 1992; Kaur and Dhawan 1994; Alam and Maughan 1995; Ganesh et al. 2000; De Boeck et al. 2004a; Das and Das 2005; Hashemi et al. 2008; Roopadevi et al. 2011); 96 h LC_50_ for Cd between 1.25 and 159 μM (Chouikhi 1979; Alkahem 1993; Suresh et al. 1993; Ramesha et al. 1996; Malekpouri and Asghar 2011; Kondera et al. 2014); and 96 h LC_50_ for Zn between 149 and 769 μM (Tishinova 1975; Hattink et al. 2006; Malekpouri and Asghar 2011).

Therefore, firstly an acute exposure experiment was setup to determine the toxicity and bioaccumulation of copper, cadmium and zinc in common carp under the standardized conditions used in our study. Subsequently, two sublethal equitoxic exposure concentrations were chosen for each metal corresponding to 50% and 10% of the calculated LC_50_-values. The main goal of this second, subacute exposure experiment was to identify the histopathological and physiological effects on the gill tissue of common carp (*Cyprinus carpio*). We hypothesized that we would find metal specific alterations, with copper interfering with sodium transport and inducing more severe gill alterations, and cadmium and zinc primarily interfering with calcium homeostasis and resulting in limited gill damage.

## Materials and Methods

### Test animals

Common carp (*Cyprinus carpio)* juveniles were obtained from the fish hatchery at Wageningen University, The Netherlands and kept in reconstituted freshwater made from deionized water (Aqualab, VWR International), supplemented with 4 salts (VWR Chemicals): NaHCO_3_ (96 mg/L), CaSO_4_.2H_2_O (60 mg/L), MgSO_4_.7H_2_O (123 mg/L), KCl (4 mg/L) to reach moderately-hard water (conductivity 308 ± 2.5 μS/cm), as defined by the US Environmental Protection Agency (Agency 2002). Each 200L tank was equipped with aeration stones and a biofilter to ensure optimal water quality. The photoperiod was set at 12 hours light, 12 hours dark, and the water temperature was maintained at 20 ± 1.5°C throughout the acclimation period and experiment. A commercial trout feed (Start Premium, 1mm, Coppens) was given *ad libitum* during the acclimation period but the fish were fastened starting from 1 day before the trial, and throughout the whole experimental period.

### Acute exposure experiment

Exposure tanks consisted of double-walled 10L polypropylene (PP) buckets, each filled with 9 L of EPA medium-hard water and containing 5 fish of 2.63 ± 0.99 g. Two replicate tanks were used per treatment and in each bucket, oxygen was provided with an air stone. In order to avoid the accumulation of ammonia and other waste products, 90% of the water was changed daily. To minimize disturbance to the fish, the perforated inner bucket was lifted from the outer bucket and the fish and 1 L of water stayed behind in the inner bucket so that the remaining 8 L of water in the outer bucket could easily be replaced after which the inner bucket was reinserted. The pH (8.21 ± 0.03) and water temperature (20 ± 1°C) were monitored daily to ensure stable experimental conditions. The tank water was spiked with an appropriate volume of the stock solutions of CdCl_2_ (Merck), CuSO_4_.5H_2_O (VWR Chemicals) and ZnCl_2_ (VWR Chemicals), to reach the desired nominal concentrations of copper (0-11 μM), cadmium (0-100 μM) and zinc (0-150 μM) respectively. Water samples were taken right before and after the water renewal, to analyze for the actual metal concentrations. Measured concentrations for each treatment are given in Table 1 and varied for cadmium between 0.00-106.04 μM, for copper between 0.02-10.78 μM, and for zinc between 0.08-129.23 μM.

**Table 1.**
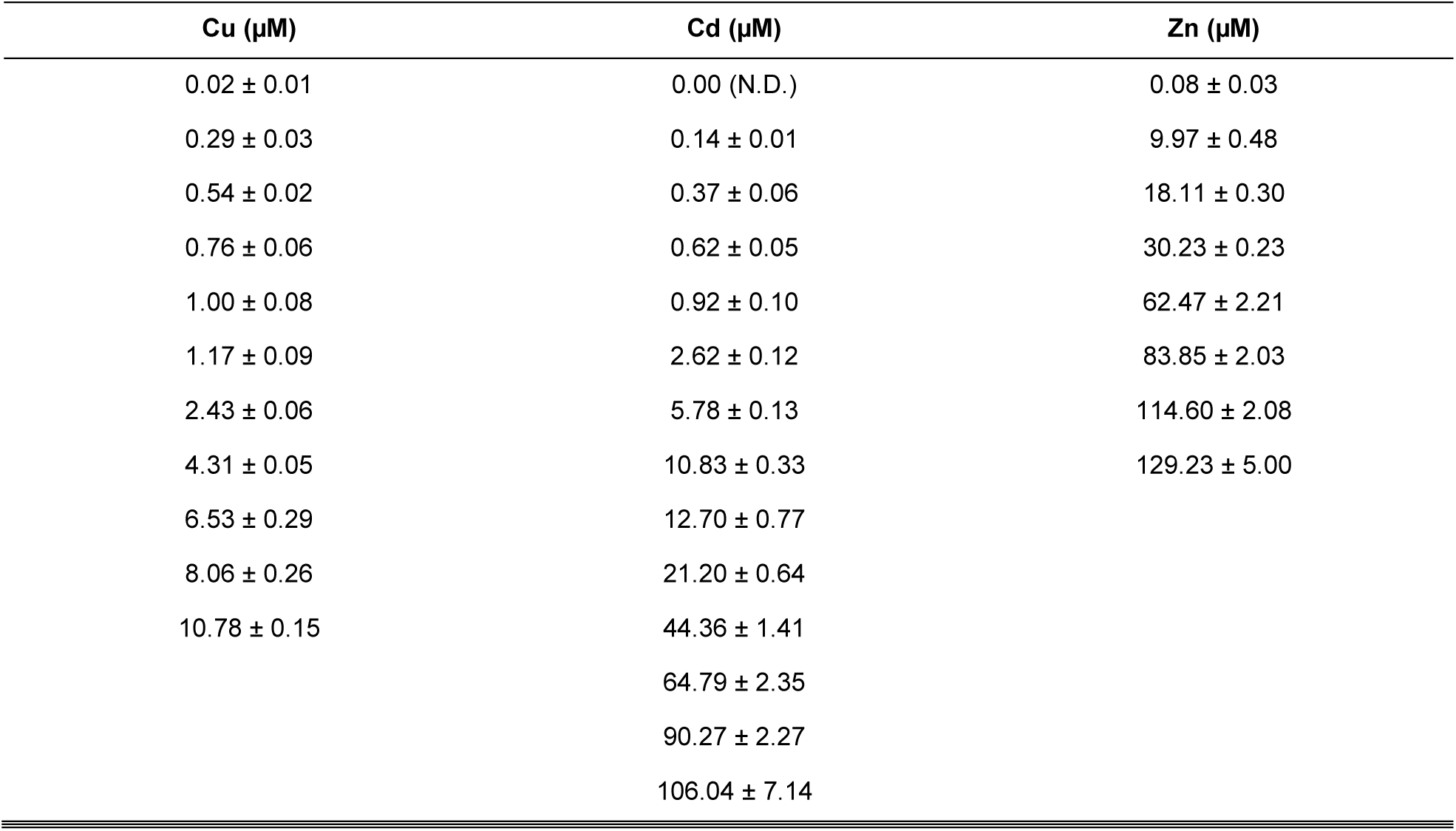
Actual metal concentrations measured in the water of the exposure tanks. The values are represented as averages (± standard error of mean) of the 2 replicate tanks, measured over the entire experimental period of 10 days (N.D. Not detectable).

Every three hours for a period of 10 days, the fish were checked for mortality or severe discomfort. Fish were considered in severe discomfort when they remained at the surface, were unable to escape to deeper water when touched, or non-responsive to touching at any water depth. When this was the case, fish were considered moribund, most likely not being able to survive the next 3 hour period, and were immediately removed and euthanized using an overdose of neutralized MS222 (Sigma). Generally, moribund fish could easily be detected in copper and zinc exposed fish, but in cadmium exposed fish death often occurred within the 3 hour intermediate period without earlier signs of severe discomfort. Dead and euthanized moribund fish were stored at −20°C for later whole-body metal analysis. After 10 days of exposure, the experiment was terminated and all remaining fish were collected, immediately euthanized with an overdose of neutralized MS222 (Sigma) and stored at −20°C for whole-body metal analysis. In total 330 fish were used of which 92 survived until the end of the experiment without any severe signs of distress, after which they were euthanized as described above. Of the remaining fish, the first 15 exposures (150 fish, 50 per metal) to concentrations based on literature data showed very quick mortality, with a 100% mortality mostly reached within 2 days, too fast to detect moribund fish before death. Of the remaining 88 fish, 31 were euthanized during the course of the experiment as they were considered in severe distress, and 57 were found dead, again, usually in the cadmium exposures.

### Sublethal metal exposures

In a similar set-up, fish from the same batch, now weighing 15.74 ± 6.67 g were stocked in the 10L PP buckets and held in an acclimated room, with a starting density of 6 fish per bucket and 3 replicate buckets per treatment. The pH and temperature was monitored daily with a field meter (Hach HQ30d) and water was renewed daily as described above. It was chosen to expose the fish to control conditions and a low and a high sublethal concentration of each metal, respectively 10% and 50% of the 96h LC_50_ obtained from the acute exposure experiment. The actual concentrations (mean ± SEM) measured in the water samples were for cadmium below detection limit for control, 0.018 ± 0.000 and 0.083 ± 0.002 μM, for copper 0.013±0.002 for control, and 0.074 ± 0.002 and 0.333 ± 0.007 μM for exposures, and for zinc 1.477±0.714 for control, 2.547 ± 0.052 and 10.765 ± 0.570 μM for exposures. Each exposure condition was executed in triplicate and lasted for 7 days. On day 1, 3 and 7, six fish were sampled from each exposure group (2 per bucket) for metal and electrolyte analysis. The fish were euthanized with an overdose of neutralized MS222 (Sigma) and thoroughly rinsed before dissection of the gills. In the final sampling, the second gill arch from the left side was collected for histological analysis.

### Determination of metal- and electrolyte content in tissue

For the acute exposure experiment, whole bodies were acid digested, whereas for the subacute exposure experiment the gill arches were collected for metal analysis. The samples were dried at 60°C for 48 hours. They were set to cool down in a desiccator before the dry weight was taken on a precision balance (Mettler AT261 DeltaRange). Next, trace-metal-grade HNO_3_ (69%, Seastar Chemicals) and H_2_O_2_ (29%, Seastar Chemicals) was added to the samples, blanks and standard reference material (SRM-2976, Mussel tissue, National Institute of Standards and Technology). The digestion process was initiated at room temperature for 12 hours, followed by a 30 minute-incubation in a hot block (Environmental Express) at 115°C. The total metal content in the digested samples was determined with a quadrupole inductively coupled plasma mass spectrometer (ICP-MS; iCAP 6000 series, Thermo Scientific). The calculated recoveries of the reference material were 102.67, 94.62 and 100.07% for cadmium, copper and zinc respectively, which indicates an adequately high accuracy of the digestion methodology.

### Histology of the gills

The left second gill arch of control and 7-day exposed fish was collected for histological analysis and placed in 4% aqueous formaldehyde (Sigma). The fixation period lasted for 48 hours, before the gills were transferred to 70% ethanol. Samples were later dehydrated in a graded ethanol series with an automatic tissue processor (Leica TP 1020), cleared in xylene, embedded in paraffin and subsequently sectioned transversely at nominal thickness of 5 μm. Slides were stained with a linear stainer (Leica ST4040) using hematoxylin and eosin (H/E) (Bancroft and Stevens, 1977). Micrographs presented in the paper were taken with a Leica DM LB microscope equipped with a Leica DFC 295 camera.

### Semi-quantitative scoring system

For the assessment of the intensity and extent of histopathological alterations in gills, a semi-quantitative scoring system developed by Bernet et al. (1999) was used. Each slide is blinded and evaluated for the extent of histopathological alterations by assigning the following score value: 0 - no alteration present; 2 - mild occurrence, 4 - moderate occurrence and 6 - severe occurrence. Intermediate values were not used, and consensual agreement between two histopathologists was done if ambiguity was present. The list of alterations found in the present study is shown in the results section (table 4), and each alteration is marked with the importance factor (w), ranging from 1 (minimal alteration) to 3 (alteration of marked importance), according to Bernet et al. (1999). Moreover, each alteration is classified in one of the following reaction patterns: circulatory (telangiectasis, hyperemia, oedema of primary epithelium and secondary epithelium), regressive (structural and architectural alterations, goblet cells in secondary lamellae, necrosis), progressive (hypertrophy, hyperplasia of epithelium and goblet cells), inflammatory (infiltration of leukocytes) and neoplastic changes (not found in the present study). For quantification of histopathological alterations, and obtaining a reaction index and a total gill index, following formulas were used:

a. Reaction index of gills (I_rp_)

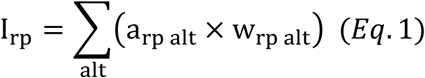
b. Histopathological index of the gills:

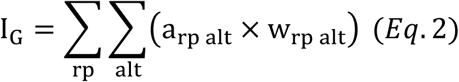

where *I*_*G*_ represents the gills index, *rp*-reaction pattern, *alt*- the alteration, *a*- score value, and *w*- importance factor.

**Table 2.** Parameters used for the logistic function to plot the dose-response relationship for the 96h and 240h exposure experiment. where “e” represents the LC50, “b” is the slope around e, and c and d are the respectively the lower and upper limit.

| Element | Duration | b | c | d | e |
| --- | --- | --- | --- | --- | --- |
| Cu | 96h | -7.99 | -0.37 | 97.16 | 0.76 |
|  | 240h | -7.69 | -0.40 | 98.62 | 0.77 |
| Cd | 96h | -1.004 | -55.36 | 97.16 | 0.20 |
|  | 240h | -14.64 | -0.17 | 98.62 | 0.22 |
| Zn | 96h | -1.80 | -22.20 | 97.16 | 29.89 |
|  | 240h | -6.03 | -0.10 | 98.62 | 26.03 |

**Table 3.** The correlation between accumulated metals and electrolyte level expressed as adjusted R-squared and corresponding p-values, N = 18.

|  |  | Cu-exposure |  | Cd-exposure |  | Zn-exposure |  |
| --- | --- | --- | --- | --- | --- | --- | --- |
|  |  | Adj R<sup>2</sup> | p-value | Adj R<sup>2</sup> | p-value | Adj R<sup>2</sup> | p-value |
| Na | Day 1 | -0.05 | 0.67 | -0.04 | 0.5 | -0.03 | 0.49 |
|  | Day 3 | **0.27** | **0.02\*** | 0.07 | 0.17 | -0.04 | 0.58 |
|  | Day 7 | **0.8** | **2.96e-07\*\*** | 0.09 | 0.16 | -0.01 | 0.38 |
| K | Day 1 | **0.27** | **0.03\*** | -0.04 | 0.53 | -0.05 | 0.68 |
|  | Day 3 | -0.05 | 0.59 | -0.04 | 0.53 | -0.04 | 0.54 |
|  | Day 7 | 0.01 | 0.3 | -0.09 | 0.98 | -0.04 | 0.59 |
| Mg | Day 1 | **0.22** | **0.03\*** | -0.06 | 0.74 | -0.06 | 0.88 |
|  | Day 3 | -0.04 | 0.52 | 0.10 | 0.14 | -0.07 | 0.94 |
|  | Day 7 | -0.04 | 0.66 | -0.05 | 0.53 | -0.04 | 0.60 |
| Ca | Day 1 | 0.15 | 0.07 | 0.00 | 0.34 | -0.05 | 0.64 |
|  | Day 3 | -0.06 | 0.74 | -0.08 | 0.93 | 0.05 | 0.20 |
|  | Day 7 | -0.04 | 0.55 | -0.08 | 0.74 | -0.04 | 0.54 |

**Table 4.**
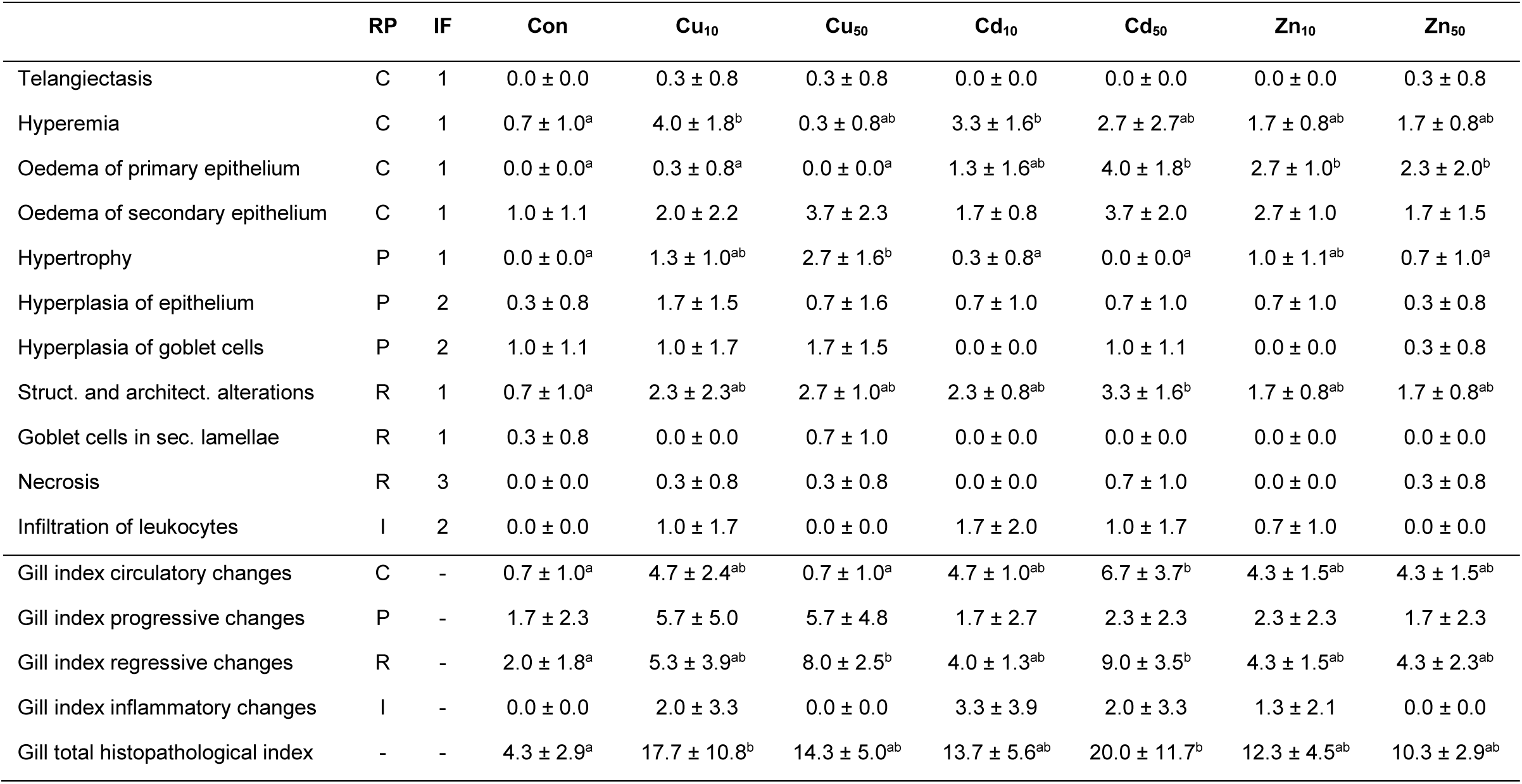
Mean values of histopathological scores of common carp from experimental groups. Scores are presented as mean values ± SD; tissue alterations were scored as follows: 0 = none, 2 = mild, 4 = moderate and 6 = severe; mean values followed by different superscript letters in the same row were significantly different (either one-way ANOVA, followed by Tukey’s HSD test or Kruskal-Wallis test, in both cases p < 0.05); IF - importance factor; RP - reaction pattern: C - circulatory changes, P - progressive changes, R - regressive changes, I - inflammatory changes.

### Statistics

GraphPad Prism 7.00 (GraphPad Software) was used to calculate the incipient lethal level (ILL) with the provided one-phase decay equation (Van Ginneken et al. 2017). All other data analysis was done with the open source software package R (version 3.4.0), and graphs were made with the *ggplot2 package* (Wickham 2009). The normality of the data and homogeneity of the variances was tested with the Shapiro-Wilks and Levene’s test, respectively. Parametric datasets were analyzed with ANOVA, followed by Tukey HSD test to make pairwise comparisons of significant differences. If the assumptions for parametric tests were not met, a Kruskal-Wallis and Dunn test were executed on the dataset. The dose-response curve (drc) package (Ritz and Streibig 2005) was used to fit a four-parameter logistic model (Equation 3) to the mortality dataset obtained from the acute toxicity test. A lack-of-fit test was executed to evaluate the acceptability of the model. Subsequently, the parameter estimates, including the LC_50_-values, and their corresponding standard deviations and p-values could be extracted from the model.

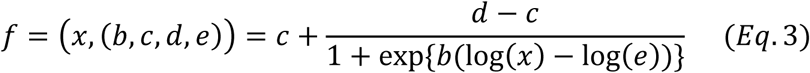

where “e” represents the LC_50_, “b” is the slope around e, and c and d are the respectively the lower and upper limit.

## Results

### Acute metal exposure

The exposure concentrations initially selected based on literature data resulted in high mortalities within the first 4 days of exposure. Therefore, additional tests were set up at lower concentrations to gather reliable toxicity data, which did end up in 11 tested concentrations for copper, 14 for cadmium and 8 for zinc (Table 1). The copper and zinc tests induced clear dose-dependent mortality, where the higher concentrations caused 100% mortality for all the fish, while the tanks spiked with lower doses of the metals, were characterized by intermediate to zero mortality (Fig. 1). In contrast, 12 out of the 14 tested concentrations for cadmium caused 90-100% mortality in the tanks, while the lowest 2 concentrations did not result in any dead fish over the entire experimental period. The fit of the four-parameter logistic model to the mortality data is presented in Figure 1, and the lack-of–fit test resulted in a p-value of 0.670 which indicates that the model is reliable to describe the dose-response relationships. Table 2 summarizes the values of the parameters describing the individual dose-response curves of the three metals, assuming both 96h- and 240h exposure scenarios. When tested over an experimental period of 10 days, zinc can be considered the least toxic metal of this study. Instead, waterborne copper and cadmium are 100-times more lethal towards common carp juveniles. The 96h LC_50_ values were calculated based on the four-parameter logistic model as well and resulted in 0.20±0.16 μM (p=0.22) for cadmium, 0.77±0.03 μM (p=2.20e^-16^) for copper, and 29.89±9.03 μM (p=1.40e^-03^) for zinc.

**Figure 1.**
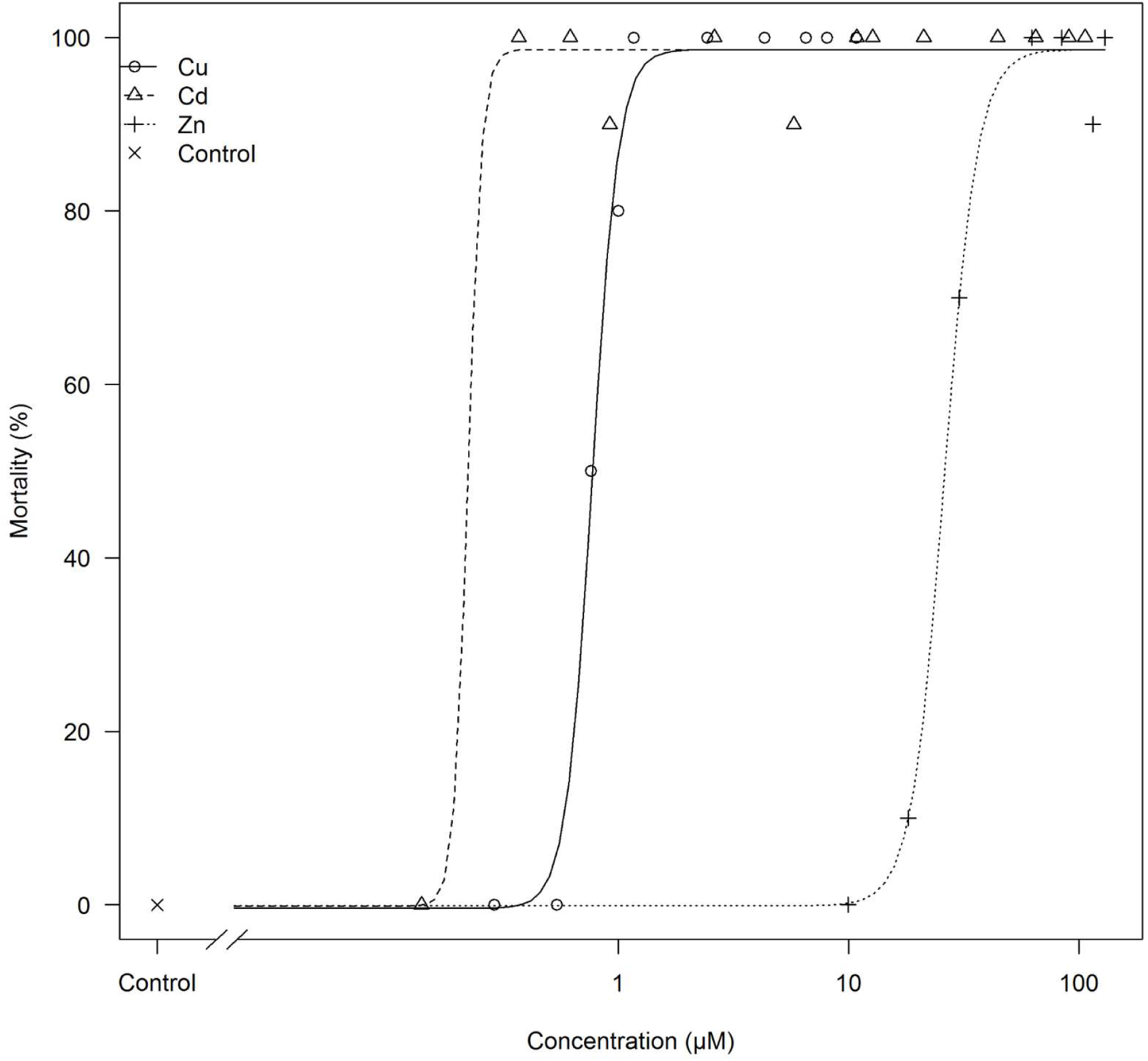
The four-parameter logistic fit of the model to the data set of copper, cadmium and zinc exposure for 240 hours. Mortality is plotted as a function of the nominal concentration of the respective metal ion (in μM).

Figure 2 shows the trend in lethal concentration of the three metals over time. The asymptotes of each curve are approaching the concentrations below which 50% of the fish will live indefinitely, i.e. the incipient lethal level (ILL). In case of a copper exposure (Fig. 2a) this level will be reached after 3 days and for cadmium (Fig. 2b) and zinc (Fig. 2c) after 5 and 6 days of exposure, respectively. The ILL for copper, cadmium and zinc calculated in this experiment are respectively 0.77 μM, 0.16 μM and 28.33 μM.

**Figure 2.**
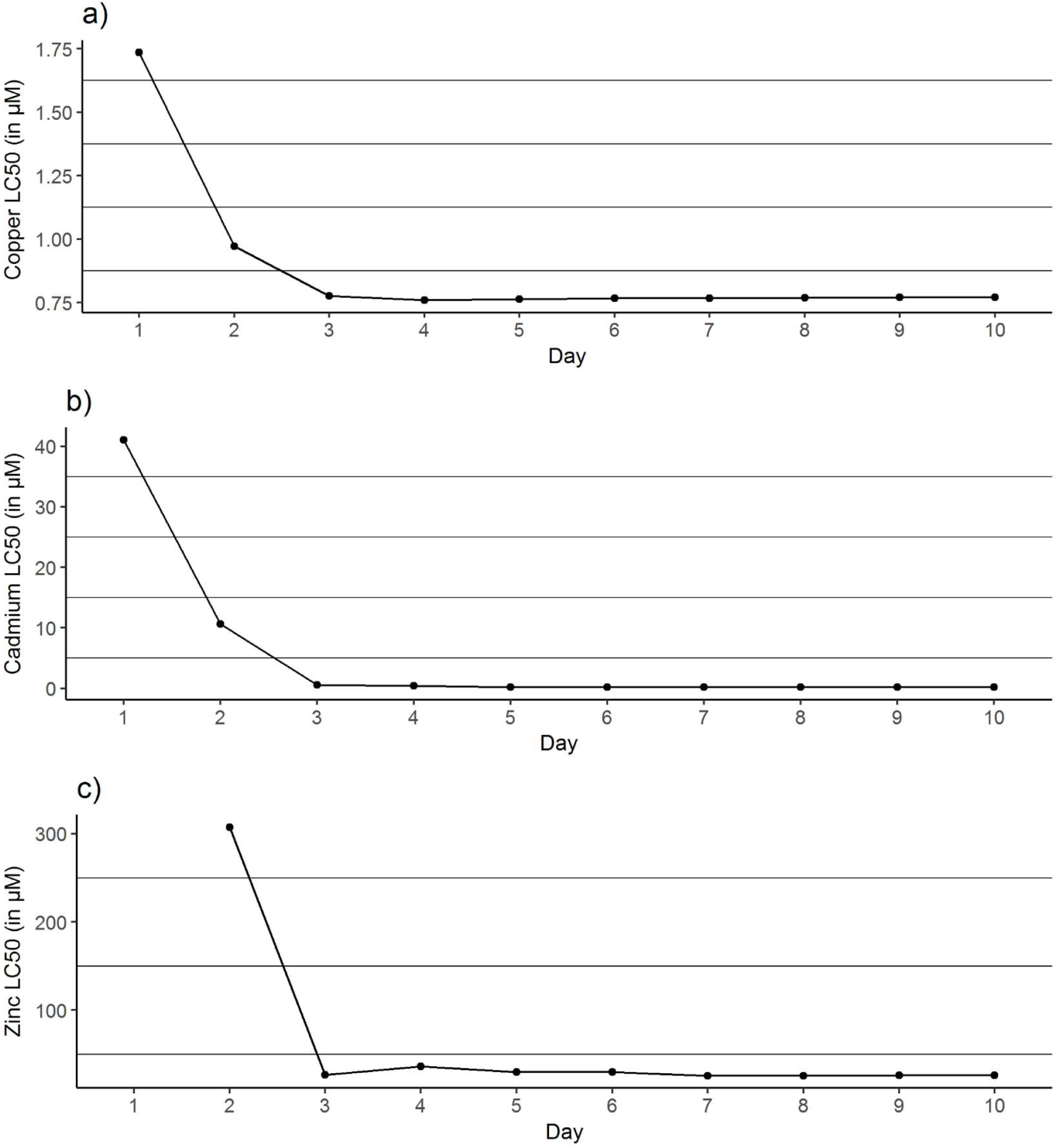
The trend in lethal level for 50% of common carp juveniles ((LC_50_) for the three metals copper (a), cadmium (b) and zinc (c) tested over a time span of 10 days (240 hours).

In a first analysis, a comparison was made between the metal content in the survivors and the non-survivors of the exposure experiment. The respective values for cadmium, copper and zinc are corrected for baseline metal content measured in the control fish, in order to get a better view on actual metal accumulation due to the exposure. In the copper exposure experiment the survivors accumulated on average 0.20 ± 0.01 μmol/g DW and the fish that died 0.17 ± 0.01 μmol/g DW. For the zinc exposures a metal content of 1.30 ± 0.27 μmol/g DW and 1.05 ± 0.27 μmol/g DW was measured in the survivors and dead fish, respectively. Neither for the copper exposure, nor for the zinc exposure these differences between survivors and victims were found to be significantly different. In contrast, there was a significantly (p=0.001) higher cadmium content measured in the fish which died after the cadmium exposure (0.26 ± 0.03 μmol/g DW) compared to the survivors (0.05 ± 0.02 μmol/g DW).

In a second analysis a concentration dependent uptake rate was calculated, where the accumulated metal content was corrected for the duration of exposure which obviously differed depending on the time of mortality. For copper exposures, copper uptake rate is very low and similar (0.0002-0.0008 μmol/g DW/h) for the 6 lowest exposure concentrations (0.2-2.2 μM). Starting from an exposure dose of 4.4 μM copper, the uptake rate starts to increase, with the highest rates observed for the fish exposed to 11 μM copper (Fig. 3a). The fish exposed to different concentrations of cadmium showed a similar pattern in uptake with a minimal increase (0.00-0.002 μmol/g DW/h) for the 9 lowest concentrations, and a notable rise in uptake rate starting from an exposure concentration of 40 μM (Fig. 3b). The fish exposed to zinc did not show such a concentration dependent increase in uptake rates (Fig. 3c). All calculated uptake rates were within the same range (0.10-0.52 μg/g DW/h), except for the fish exposed to 10 μM (0.002 μmol/g DW/h) and 20 μM (0.004 μmol/g DW/h) of zinc.

**Figure 3.**
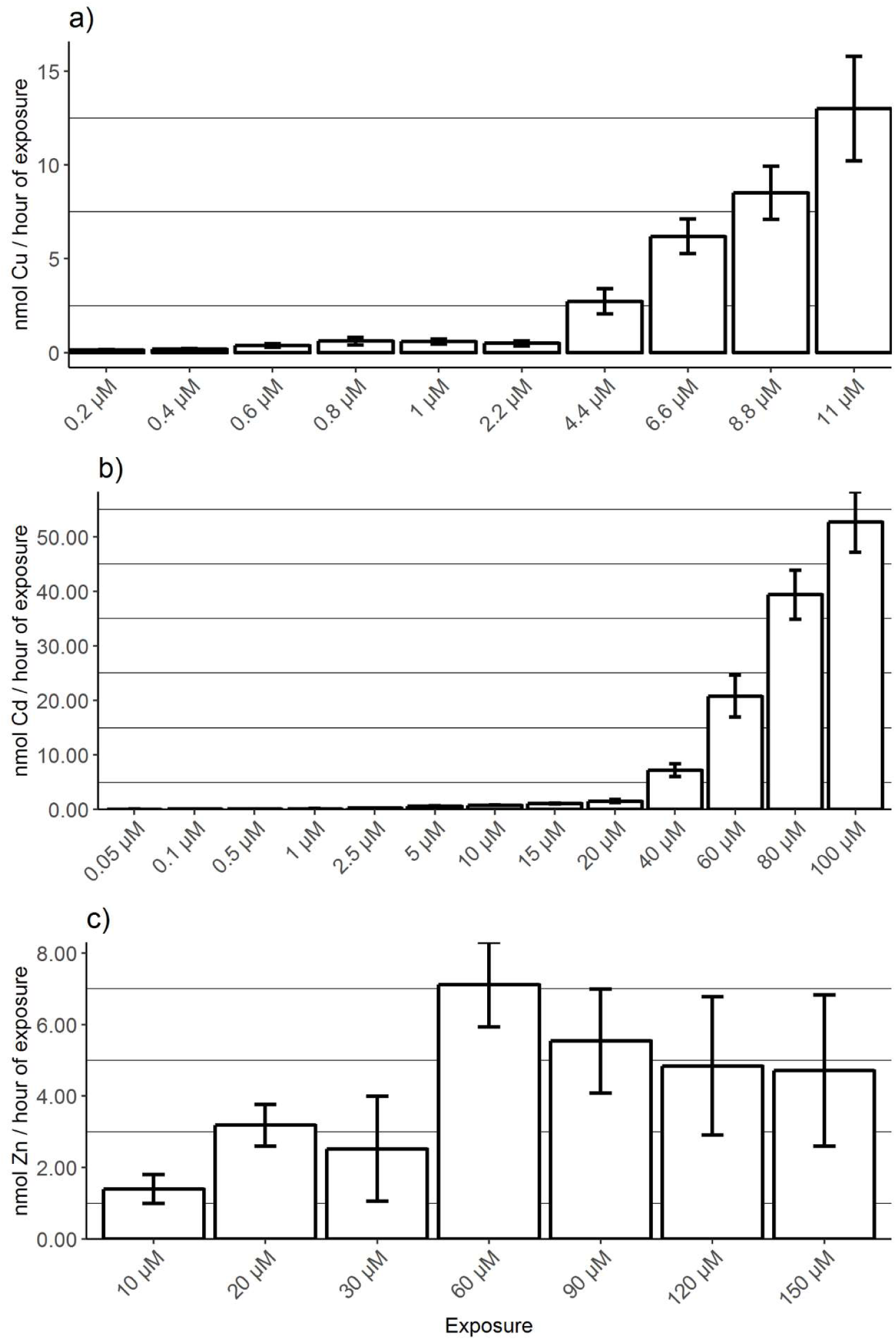
Metal uptake rate (mean ± SE) during the 240 hours toxicity screening test for the different exposure concentrations (in μM), for copper (a), cadmium (b) and zinc (c) exposure. The total metal content (in μg/g dry weight) measured in the whole body is corrected for the average value of the respective metal in the control fish. The y-axis expresses the uptake rate in hours of exposure with error bars as standard error of mean.

### Sublethal metal exposure

At the start of the experiment cadmium was not detectable in the fish gills, whereas the copper and zinc content was respectively 0.08 ± 0.006 μmol/g and 16.45 ± 1.19 μmol/g dry weight. A time- and concentration-dependent increase in copper accumulation in the gill tissues was seen as a result of waterborne copper exposure (Fig. 4a). At day 1 and 3 only the fish exposed to the high copper concentration (50% of 96h LC_50_) did accumulate significant higher levels of copper than the control fish. After 7 days, both groups exposed to a low and high copper concentration had significantly more copper in their gill tissue compared to the control group (0.23 ± 0.01 μmol/g and 0.66 ± 0.05 μmol/g, respectively). As for the whole-body metal analysis, the actual accumulation of copper can also be estimated by correcting the measured level with the average value of the control fish. Such a calculation shows that the fish exposed to the low and high dose of copper accumulated approximately 0.13 and 0.56 μmol copper. This corresponds respectively to a 123 and 538% increase in gill copper levels after 7 days of exposure and a net uptake rate of 0.0008 and 0.003 μg/g DW/h respectively.

**Figure 4.**
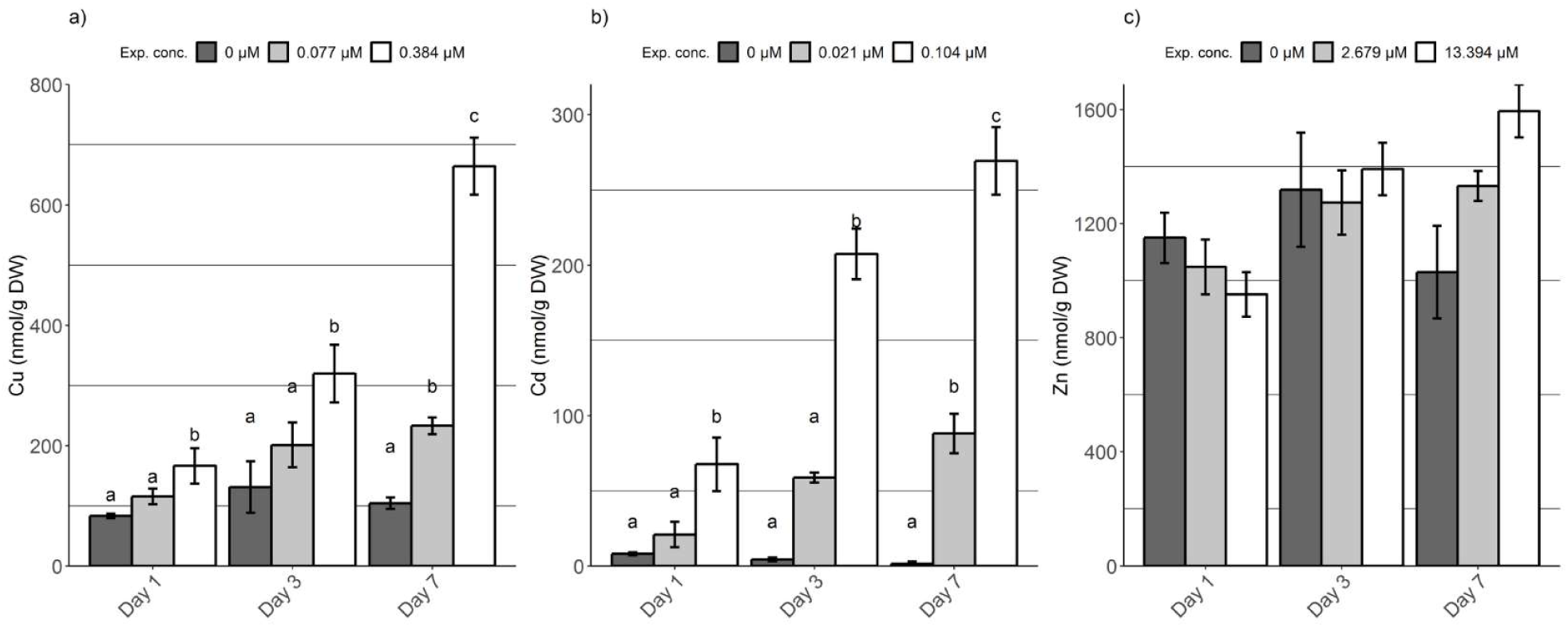
Total copper (a), cadmium (b) and zinc (c) accumulated in the gills expressed in micrograms per gram of dry weight (mean ± SE). The metal content was measured after 1, 3 and 7 days of exposure for the 3 experimental groups; control (dark grey), low dose (light grey) and high dose (white). Day 0 refers to the time zero control group (black). Letters denote significant differences among treatments and within 1 sampling day, found with Dunn’s test of multiple comparisons (Bonferroni adjusted p-value rejects H_0_ if p<=0.05/2).

Cadmium levels in the gills did also increase in a time- and concentration-dependent manner (Fig. 4b). At day 1, the fish exposed to the high dose of cadmium did accumulate significantly more cadmium than the control fish, and this remained the case for the entire experiment. A similar pattern was seen for the treatment group exposed to a low dose of cadmium, however statistically significant higher levels of cadmium in the gill tissue compared to the control group were only observed after 7 days of exposure. At the end of the experiment, low- and high dose treated groups accumulated on average 0.09 ± 0.01 and 0.27 ± 0.02 μmol cadmium per gram of dry gill tissue which corresponds to a net uptake rate of 0.0005 and 0.001 μmol/g DW/h respectively.

The variation in zinc content of the gills of the carp juveniles was rather high during the entire experiment (Fig. 4c), also for the fish which were not exposed to zinc. No significant differences in zinc accumulation could be observed as a result of the treatment. It is only at day 7 that the expected trend starts to take shape with small, but dose dependent differences in zinc accumulation among treatment groups. The fish exposed to the lowest and highest dose had respectively 20.36 ± 0.81 and 24.39 ± 1.42 μmol/g Zn in their gill tissue on the last sampling day. When corrected for the zinc content in the control fish, this resulted in a net accumulation of 4.61 and 8.63 μmol zinc or a 30 and 54% increase, respectively. This corresponds to an uptake rate of 0.03 and 0.05 μg/g DW/h respectively.

A linear regression model was used to fit the specific electrolyte levels (Na, K, Mg and Ca) to the accumulated metals (Table 3). Copper was the only metal which induced a significant change on the electrolyte content in the gill tissue. On day 1 a transient slightly positive correlation with K and Mg levels (resp., R^2^ = 0.27, p=0.02 and R^2^=0.22, p=0.03) with respect to accumulated copper was observed. Gills’ sodium levels are clearly affected by the copper content. Starting from day 3, lower sodium contents are significantly associated with increasing copper concentrations (R^2^=0.27, p=0.02). This trend continues on day 7 where a highly significant correlation between sodium and copper was observed (R^2^=0.8, p=2.96e^-7^). Neither the zinc-exposure, nor the cadmium exposure did reveal any significant correlation between accumulated metals and gill electrolyte levels.

Histopathological alterations found in gills of common carp (Table 4) were generally mild at these sublethal exposure levels and no dose-dependent relationship could be established using semiquantitative scoring. At the end of experiment, histopathological changes in the control group were either totally absent or some of them were rarely noticed (Fig. 5a). Moderate scores were present only in alterations with low histopathological significance, according to importance factor. They include hyperemia (Fig. 5b) in fish from Cu_10_ and Cd_10_ groups, hypertrophy (Fig. 5c) in Cu_50_ group, structural and architectural alterations (Fig. 5d) in Cd_50_ group, and oedema of primary epithelium (Fig. 5e) in Cd_50_, Zn_10_ and Zn_50_ groups. As we defined moderate scores ranging from 2 to 4 in the exposed carp, the majority of them were significantly higher compared to the control group. Gill necrosis, as an irreversible alteration with highest importance factor, was present only in small scattered areas. (Fig. 5f).

**Figure 5.**
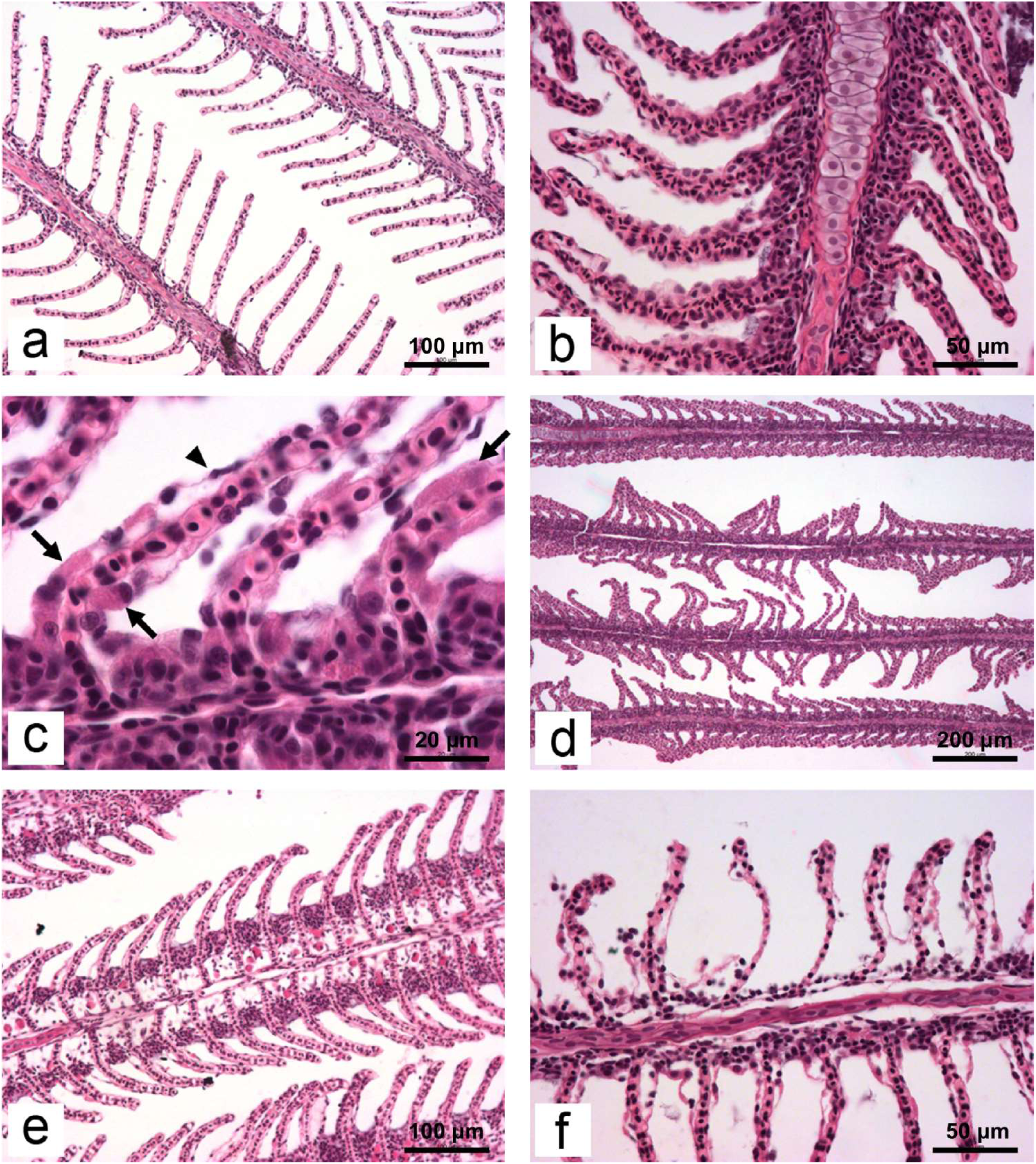
Micrographs of some of histopathological alterations found in the gills of common carp in the present study: a) normal tissue structure (H/E x200); b) note the extensive hyperemia on secondary lamellae at the left side of primary filament (H/E x400); c) hypertrophy of squamous epithelial cells (arrows); compare thickness of these cells to non-pathological cell (arrowhead). Furthermore, note the size of the nuclei in both type of cells (H/E x1000); d) disturbed architecture of secondary lamellae: curling and fusion (H/E x100); e) oedema of primary filament (H/E x200); f) total rupture of epithelium leading to necrosis (H/E x400).

Calculated HP indices of different reaction patterns observed after 7 days of exposure are presented in Table 4. As with the individual alterations, the control group showed lowest values in all indices. In the HP index of progressive changes (I_GP_) and in the HP index of inflammatory changes (I_GI_), no statistical significance was established between groups. In the HP index of circulatory changes (I_GC_), group Cd_50_ was higher compared to both control and Cu_50_ group (p<0.05), while in the HP index of regressive changes (I_GR_), fish exposed to highest concentrations of Cu and Cd had elevated score values compared to the control (P<0.05). The cumulative index of all histopathological alterations, I_GT_, showed that fish in the Cu_10_ and Cd_50_ groups underwent the highest impact compared to fish from control group (p<0.05).

## Discussion

The acute exposure experiment indicated a much higher toxicity of copper and cadmium, compared to what was expected based on the literature review. Several explanations for this outcome can be given. First of all, the size and age of the fish will greatly influence the tolerance of fish towards a pollutant (Kousar and Javed 2012). De Boeck et al. (2004), conducted a similar experiment with copper-exposed common carp and found a 96h LC_50_-value of 10.4 μM. The carp used for that experiment had an average weight of 60.6 ± 27.8 g, which most likely explains the over 10-fold higher tolerance to waterborne copper, compared to the 2.6 g juveniles used for this experiment. For cadmium and zinc also higher lethal levels were reported, which can similarly be attributed to the fact that larger fish were used (Naji et al. 2007; Kondera et al. 2014). Secondly, the water chemistry does also play a major role in metal availability, and thus toxicity. Calcium and sodium ions are typically decreasing the toxicity of metals at the biotic ligand, the fish gills, and are also known for reducing the levels of histopathological alterations when fish are exposed to metals in the hard water (Pratap and Wendelaar Bonga 1993; Liu et al. 2010; Crémazy et al. 2017). Calcium is also known for its protective role, due to the change in gill permeability or competition with other ions for the binding sites (Hogstrand et al. 1994; Hollis et al. 2000). The use of medium-hard water, in which fish were exposed during the present study, could explain lower number of histopathological alterations compared to other studies and the generally mild intensity of changes. Not all researchers do consistently report the chemical characteristics of the exposure water, which makes it hard to relate differences in experimental outcome to this aspect. A study by Hattink et al. (2008) does provide detailed information on the water chemistry. The carp were comparable in size to the fish used here, but the obtained LC_50_-value of zinc was approximately 5 times higher (149 μM) than in our experiment. This higher tolerance towards zinc found by the researchers is probably related to the nearly 6-fold higher concentration of Ca^2+^-ions in the exposure water (Hattink et al. 2006). Further, physical parameters like pH and temperature are also known to influence the bioavailability and uptake rate of metals, which might contribute to differences in lethal concentrations with previous studies (Blust 1988; Furuta et al. 2007).

In toxicology, a 96-hour exposure experiment is the most common procedure to evaluate the toxicity of a particular compound. Yet, this time span has been debated within the scientific community, as some studies show that the concentration below which 50% of the fish will survive, cannot be determined after only 4 days of exposure (Hasan and Macintosh 1986; De Boeck et al. 2004b). In the present case it was seen that the time to reach the ILL is pollutant-dependent. The results suggest that in our copper experiment, the classic 4-day exposure period was long enough to provide reliable toxicity data. However, incipient lethal levels of cadmium and zinc were only reached after 5 and 6 days of exposure, which advocates for longer exposure experiments.

The total metal accumulation was both quantified in the whole body, in the first experiment, and in the gills in the subacute exposures. In general, a clear increase in copper and cadmium content, proportionally to the treatment dose was observed in both experiments. Zinc, however, did not show such a clear dose-dependent increase. It can be postulated that this slower accumulation of zinc is related to a combination of the relatively low waterborne zinc concentrations compared to the calcium concentration. As mentioned before, the latter can competitively inhibit the uptake of zinc (Spry and Wood 1989). However, this is even more true for cadmium, where the dose-dependent uptake was obvious. More likely, the naturally high zinc content in the gill tissue (16.45 μmol/g DW) demands a longer exposure period to discern between control and treatment groups. Finally, the fact that zinc is an essential metal implies that fish are able to control the overall uptake of this metal respective to their own needs. Several transmembrane proteins responsible for the import of zinc, have been identified for fish and were found to have a different affinity for waterborne zinc (Qiu and Hogstrand 2005). Previous studies suggest that the activity of these high-affinity zinc importers can be regulated as a consequence of environmental zinc concentrations (Hogstrand et al. 1998; Alsop and Wood 2000), so this might explain the slow increase in zinc content in the gills. As for zinc, copper is also essential for basic metabolic processes in animals. Therefore, similar regulatory mechanisms are to be expected to maintain viable levels in the cells. Clearly, carp were less able to control copper accumulation when compared to zinc accumulation at equitoxic concentrations. Three uptake pathways for copper have been suggested in fresh water fish; the apical Na^+^-channel, a divalent cation transporter (DMT1) and a high affinity copper transporter (Ctr1) (Grosell and Wood 2002; Bury et al. 2003; Mackenzie et al. 2004). From a toxicological point of view clear copper accumulation could be attributed to the failure of regulating one or some of the above-mentioned uptake pathways.

When exposed to Cu, we found some small scattered areas of necrosis as reported before (Cerqueira and Fernandes 2002; Al-Bairuty et al. 2013). More common was partial lamellar fusion in the form of fused tips of secondary lamellae. Partial or even complete lamellar fusion of lamellae has been detected before under copper exposure, mostly at much higher Cu concentrations (Karan et al. 1998; Muhvich et al. 1995), and could compromise respiration.

Hypertrophy of secondary lamellae and their tips occurred in the Cu_50_ group similar to what was seen in common carp exposed to 1.72 μM Cu (De Boeck et al. 2007), and the Cu_50_ group showed the highest level of semiquantitative scores. However, the Cu_10_ displayed higher levels of hyperemia, a consequence of an increase in blood flow caused by vasodilation of gill arteries, leading to a higher gill index. This physiological adaptation is characteristic for the second phase of the stress response (Harper and Wolf 2009) and could point out to an adaptation phase in fish exposed to lower levels of copper. It is widely accepted that copper has a damage-repair mode of action and that deleterious effects are the most pronounced from 4 to 24 h following exposure (De Boeck et al. 2001; De Boeck et al. 2007) after which gills are starting to recover although full recovery is possible only in copper-free water (Cerqueira and Fernandes 2002). These histological changes in the fish gills due to copper exposure are mainly explained by the disruptive physiological effect of the element on the overall ion balance. After exposing fish to even low concentrations of waterborne copper, a drop in sodium and chloride levels is occurring in the plasma as was seen here as well, and at the same time an increase in potassium levels is observed. This effect was seen in a number of freshwater fish, such as common carp, Nile tilapia (*Oreochromis niloticus*) and *Prochilodus scrofa* (Cerqueira and Fernandes 2002; De Boeck et al. 2004b; Monteiro et al. 2005). The main explanation of this sodium effect lies within shared use of sodium and copper ions of the apical sodium channel causing a competitive inhibition of sodium uptake (Grosell and Wood 2002) and the negative correlation between the level of copper in gills and Na^+^/K^+^-ATPase activity (De Boeck et al. 2001; Monteiro et al. 2005). Hyperplasia of mitochondria rich cells, rich in Na^+^/K^+^-ATPase, as well as mucous cells occurs as a compensatory mechanism for the loss of ion uptake capacity (Pratap and Wendelaar Bonga 1993; Grosell and Wood 2002; Monteiro et al. 2009)

Although cadmium has repetitively demonstrated to inhibit ion-transporting enzymes like Ca^2+^-ATPase and Na^+^/K^+^-ATPase (Pratap and Wendelaar Bonga 1993; Mcgeer et al. 2000; Matsuo et al. 2005) with additional competition of cadmium with Ca and K at the apical calcium channel (Shahsavarani et al. 2006), our study did not indicate any loss of these ions after an exposure to 0.021 or 0.104 μM Cd. Previous studies showed blood congestion, various degrees of lamellar hyperplasia, including complete fusion of lamellae, as well as hypertrophy and hyperplasia of chloride and mucous cells in white seabass (*Lates calcarifer*) and Nile tilapia (Thophon et al. 2003; Garcia-Santos et al. 2005). In the present study, at much lower exposure concentrations, and similarly to copper, hyperemia of secondary lamellae was significantly higher in the group exposed to the lower concentration. Ultrastructural studies showed that cadmium is responsible for enlargement of microridges in epithelial cells, but it also has an influence on chloride cells density and their apical area (Wong and Wong 2000).

Contrary to copper and cadmium, studies of zinc toxicity to fish gills and their morphological changes remained scarce. The exposure concentrations for zinc are relatively low in comparison to the high background levels in the gill tissue. This, in combination with the short exposure time not allowing zinc to accumulate up to statistically different levels among treatment groups, explains the low degree of histological effects. Nevertheless, in both zinc groups, oedema of the primary epithelium was significantly higher comparing to control group as was also found in a study where *Astyanax* aff. *bimaculatus* was exposed to higher concentrations of zinc (Santos et al. 2012). This is most likely a consequence of disrupted calcium homeostasis (Spry and Wood 1985) as a direct consequence of competition at the uptake site, the epithelial calcium channel, or because of influence of zinc on the cellular Ca^2+^ signaling (Hogstrand et al. 1995; Haase and Maret 2005; Shahsavarani and Perry 2006), although we did not measure significant drops in gill calcium levels here.

Two alterations that contributed the most to the highest levels of the total gill index score in our study are oedema and structural and architectural alterations. Both changes are not of high histopathological significance, and could easily be reverted when the stressor is removed from the aquatic environment (Poleksic and Mitrovic-Tutundzic 1994). Concerning the specificity of histopathological changes, oedema of the primary epithelium was the only alteration absent in the control and both groups exposed to copper, but showed significantly higher scores in fish exposed to Cd_50_, Zn_10_ and Zn_50_. Lamellar oedema is originating as a consequence of ultrafiltration subjected to increased arterial blood pressure, and Perry et al. (1984) compared this mechanism to glomerular filtration in the kidney. The described process is easily reversible and if stressor is removed from the water, it will happen fairly rapid (Roberts and Ellis 2012). Cadmium and zinc share common mechanisms of toxicity (Wood 2001) and this is probably the explanation for the absence of oedema in the copper exposed fish. Fish exposed to higher concentrations of copper and cadmium were exhibiting increased scores in regressive changes. Regressive changes are defined as “malformation or dysfunction of cellular structures as a result of cell damage” (Bernet et al. 1999) and this pattern is generally linked to a decreasing number of cells in the tissue, contrary to progressive changes.

## Conclusion

In conclusion, concentrations used in the present study are considerably lower compared to other acute and subacute studies mentioned in the discussion. This is related to both life-history traits of the animals, as well as the difference in environmental parameters. Yet, the exposed concentrations were much closer to the actual concentrations found in surface waters, making the study far more relevant for comparison with environmental data. There was no significant difference in metal accumulation between dead and surviving fish for copper and zinc, but a clear difference was seen for cadmium where dead fish had accumulated significantly more cadmium. Bioaccumulation patterns at equitoxic concentration differed between metals, with clear dose dependent accumulation for copper and cadmium, but not for zinc. Overall, physiological and histopathological changes at 10 and 50% of the LC50 value were mild. Metal specific alterations were reduced sodium levels in gills of copper exposed fish and oedema of the primary epithelium in zinc, and to a lesser extent cadmium, exposed fish.

## Acknowledgment

This work was supported by a TOP-BOF project [grant number 32252] to GDB, LB and RB by the University of Antwerp Research Council and by the Ministry of Education, Science and Technological Development of Serbia to BR [grant number TR31075]. The funders had no role in study design, data collection and analysis, decision to publish, or preparation of the manuscript. Further, we thank our lab technician S. Joosen for his support concerning the metal analyses.

## Compliance with Ethical Standards

The fish husbandry and experiments complied with the regulation of the Federation of European Laboratory Animal Science Associations and were approved by the local ethics committee, University of Antwerp (Permit Number: 2015-94 Project 32252). VD, LB, RB and GDB all obtained the certificate of Experimenter Category C as defined in Art 11§4 and annex 8 of the Royal Decree of April 6, 2010 concerning the protection of laboratory animals. BR (gill microscopy) and MSS (metal and electrolyte analysis) analyzed samples after the exposures were done.

All authors of this manuscript declare that there are no conflicts of interest to disclose.

## References

Agency USEP. 2002. Methods for Measuring the Acute Toxicity of Effluents and Receiving Waters to Freshwater and Marine Organisms. Fifth Edit. Washington, DC 20460.

Al-Bairuty GA, Shaw BJ, Handy RD, Henry TB. 2013. Histopathological effects of waterborne copper nanoparticles and copper sulphate on the organs of rainbow trout (Oncorhynchus mykiss). Aquat Toxicol. 126:104–115.

Alam MK, Maughan OE. 1992. The Effect of Malathion, Diazinon, and Various Concentrations of Zinc, Copper, Nickel, Lead, Iron, and Mercury on Fish. Biol Trace Elem Res. 34:225–236.

Alam MK, Maughan OE. 1995. Acute toxicity of heavy metals to common carp (Cyprinus carpio). J Environ Sci Heal. 30(8):1807–1816.

Alkahem HF. 1993. Ethological Responses and Changes in Hemoglobin and Glycogen Content of the Common Carp, Cyprinus carpio, Exposed to Cadmium. Asian Fish Sci Metro Manila. 6(1):81–90.

Almeida JA, Diniz YS, Marques SFG, Faine LA, Ribas BO, Burneiko RC, Novelli ELB. 2002. The use of the oxidative stress responses as biomarkers in Nile tilapia (Oreochromis niloticus) exposed to in vivo cadmium contamination. Environ Int. 27(8):673–679.

Alsop DH, Wood CM. 2000. Kinetic analysis of zinc accumulation in the gills of juvenile rainbow trout: Effects of zinc acclimation and implications for biotic ligand modeling. Environ Toxicol Chem. 19(7):1911–1918.

Bernet D, Schmidt H, Meier W, Wahli T. 1999. Histopathology in fish proposal for a protocol to assess aquatic pollution. J Fish Dis. 22:25–34.

Blust R. 1988. Evaluation of microwave heating digestion and graphite furnace atomic absorption spectrometry with continuum source background correction for the determination of iron, copper and cadmium in brine shrimp. J Anal At Spectrom. 3(2).

De Boeck G, Meeus W, De Coen W, Blust R. 2004a. Tissue-specific Cu bioaccumulation patterns and differences in sensitivity to waterborne Cu in three freshwater fish: rainbow trout (Oncorhynchus mykiss), common carp (Cyprinus carpio), and gibel carp (Carassius auratus gibelio). Aquat Toxicol. 70:179–188.

De Boeck G, Meeus W, De Coen W, Blust R. 2004b. Tissue-specific Cu bioaccumulation patterns and differences in sensitivity to waterborne Cu in three freshwater fish: Rainbow trout (Oncorhynchus mykiss), common carp (Cyprinus carpio), and gibel carp (Carassius auratus gibelio). Aquat Toxicol. 70:179–188.

De Boeck G, Van der Ven K, Meeus W, Blust R. 2007. Sublethal copper exposure induces respiratory stress in common and gibel carp but not in rainbow trout. Comp Biochem Physiol - C Toxicol Pharmacol. 144(4):380–390.

De Boeck G, Vlaeminck A, Balm PH, Lock R a, De Wachter B, Blust R. 2001. Morphological and metabolic changes in common carp, Cyprinus carpio, during short-term copper exposure: interactions between Cu2+ and plasma cortisol elevation. Environ Toxicol Chem. 20(2):374–81.

Bury NR, Walker PA, Glover CN. 2003. Nutritive metal uptake in teleost fish. J Exp Biol. 206:11–23.

Casanova FM, Honda RT, Ferreira-nozawa MS. 2013. Effects of Copper and Cadmium Exposure on mRNA Expression of Catalase, Glutamine Synthetase, Cytochrome P450 and Heat Shock Protein 70 in Tambaqui Fish (Colossoma Macropomum). Gene Expr to Genet Genomics.:1.

Cerqueira CCC, Fernandes MN. 2002. Gill Tissue Recovery after Copper Exposure and Blood Parameter Responses in the Tropical Fish Prochilodus scrofa. Ecotoxicol Environ Saf. 52(2):83–91.

Chouikhi A. 1979. Choice and set up of the food chains in freshwater in order to show the bioaccumulation character of a pollutant. In: OECD-IRCHA Universite Paris-Sud, Unite d’Enseignement et de Recherche d’Hygiene et Protection de l’Homme et de son Environnement (FRE). Commission E. 2008. DIRECTIVE 2008/105/EC.

Crémazy A, Wood CM, Ng TY-T, Smith DS, Chowdhury MJ. 2017. Experimentally derived acute and chronic copper Biotic Ligand Models for rainbow trout. Aquat Toxicol. 192:224–240.

Das BK, Das N. 2005. Impacts of quicklime (CaO) on the toxicity of copper (CuSO 4, 5H 2O) to fish and fish food organisms.

Deshmukh SS, Marathe VB. 1980. Size related toxicity of copper & mercury to Lebistes reticulatus (Peter), Labeo rohita (Ham.) & Cyprinus carpio Linn. Indian J Exp Biol. 18(14):421–423.

Ebrahimpour M, Alipour H, Rakhshah S. 2010. Influence of water hardness on acute toxicity of copper and zinc on fish. Sage. 26(6).

Eyckmans M, Celis N, Horemans N, Blust R, De Boeck G. (2011). Exposure to waterborne copper reveals differences in oxidative stress response in three freshwater fish species. Aquatic Toxicology 103, 112–120.

EPA. 2016. Ecotox database. [accessed 2016 Feb 10]. https://cfpub.epa.gov/ecotox/.

Fonseca AR, Sanches Fernandes LF, Fontainhas-Fernandes A, Monteiro SM, Pacheco FAL. 2016. From catchment to fish: Impact of anthropogenic pressures on gill histopathology. Sci Total Environ. 550(February):972–986.

Fonseca AR, Sanches Fernandes LF, Fontainhas-Fernandes A, Monteiro SM, Pacheco FAL. 2017. The impact of freshwater metal concentrations on the severity of histopathological changes in fish gills: A statistical perspective. Sci Total Environ. 599–600(May):217–226.

Furuta T, Iwata N, Kikuchi K. 2007. Effects of fish size and water temperature on the acute toxicity of boron to Japanese flounder Paralichthys olivaceus and red sea bream Pagrus major. Fish Sci. 73:356–363.

Ganesh N, Gupta TRC, Katti RJ, Udupa KS. 2000. Acute toxicity of copper on three life stages of common carp, Cyprinus carpio Var. communis. Pollut Res. 19(1):91–93.

Garcia-Santos S, Fontainhas-Fernandes A, Wilson JM. 2005. Cadmium Tolerance in the Nile Tilapia (Oreochromis niloticus) Following Acute Exposure: Assessment of Some Ionoregulatory Parameters. Wiley Intersci.

Van Ginneken M, Blust R, Bervoets L. 2017. How lethal concentration changes over time: Toxicity of cadmium, copper, and lead to the freshwater isopod Asellus aquaticus. Environ Toxicol Chem. 36(10):2849–2854.

Green WW, Mirza RS, Wood CM, Pyle GG. 2010. Copper Binding Dynamics and Olfactory Impairment in Fathead Minnows (Pimephales promelas). Environ Sci Technol. 44:1431–1437.

Grosell M, Wood CM. 2002. Copper uptake across rainbow trout gills: mechanisms of apical entry. J Exp Biol. 205(Pt 8):1179–1188.

Guo Z, Ye H, Xiao J, Hogstrand C, Zhang L. 2018. Biokinetic Modeling of Cd Bioaccumulation from Water, Diet and Sediment in a Marine Benthic Goby: A Triple Stable Isotope Tracing Technique. Environ Sci Technol. 52:8429–8437.

Haase H, Maret W. 2005. Trace Elements Trace Elements Fluctuations of cellular, available zinc modulate insulin signaling via inhibition of protein tyrosine phosphatases. Med Biol J Trace Elem Med Biol. 19:37–42.

Harper C, Wolf JC. 2009. Morphologic Effects of the Stress Response in Fish. ILAR J. 50(4):387–397.

Hasan MR, Macintosh DJ. 1986. Acute toxicity of ammonia to common carp Fry. Aquaculture. 54:97–107.

Hashemi S, Blust R, De Boeck G. 2008. The effect of starving and feeding on copper toxicity and uptake in Cu acclimated and non-acclimated carp. Aquat Toxicol. 86:142–147.

Hattink J, De Boeck G, Blust R. 2006. Toxicity, accumulation, and retention of zinc by carp under normoxic and hypoxic conditions. Environ Toxicol Chem. 25(1):87–96.

Hogstrand C, Reid S, Wood C. 1995. Ca2+ versus Zn2+ transport in the gills of freshwater rainbow trout and the cost of adaptation to waterborne Zn2+. J Exp Biol. 198(Pt 2):337–348.

Hogstrand C, Webb N, Wood CM. 1998. Covariation in Regulation of Affinity For Branchial Zinc and Calcium Uptake in Freshwater Rainbow Trout. J Exp Biol. 201:1809–1815.

Hogstrand C, Wilson RW, Polgar D, Wood CM. 1994. Effects of Zinc on the Kinetics of Branchial Calcium Uptake in Freshwater Rainbow Trout During Adaptation to Waterbone Zinc. J Exp Biol. 186:55–73.

Hollis L, McGeer JC, McDonald DG, Wood CM. 2000. Protective effects of calcium against chronic waterborne cadmium exposure to juvenile rainbow trout. Environ Toxicol Chem. 19:2725–2734.

Kamunde C, Clayton C, Wood CM. 2002. Waterborne vs. dietary copper uptake in rainbow trout and the effects of previous waterborne copper exposure. Am J Physiol Regul Integr Comp Physiol. 283:69–78.

Karan V, Vitorovic S, Tutundzic V, Poleksic V. 1998. Functional enzymes activity and gill histology of carp after copper sulfate exposure and recovery. Ecotoxicol Environ Saf. 40(1–2):49–55.

Kaur K, Dhawan A. 1994. Metal toxicity to different life stages of Cyprinus carpio Linn. Indian J Ecol. 21(2):93–95.

Kondera E, Ługowska K, Sarnowski P. 2014. High affinity of cadmium and copper to head kidney of common carp (Cyprinus carpio L.). Fish Physiol Biochem. 40:9–22.

Kostic J, Kolarevic S, Kracun-Kolarevic M, Aborgiba M, Gacic Z, Paunovic M, Višnjic-Jeftic Ž, Raškovic B, Poleksic V, Lenhardt M, et al. 2017. The impact of multiple stressors on the biomarkers response in gills and liver of freshwater breams during different seasons. Sci Total Environ. 601–602:1670–1681.

Kousar S, Javed M. 2012. Evaluation of Acute Toxicity of Copper to Four Fresh Water Fish Species. Int J Agric Biol. 14(5):801–804.

Li J, Lock RAC, Klaren PHM, Swartz HGP, Schuurmans Stekhoven FMAH, Wendelaar Bonga SE, Flik G, 1996. Kinetics of Cu2+ inhibition of Na+/K+-ATPase. Toxicol. Lett. 87, 31–38.

Liu XJ, Luo Z, Xiong BX, Liu X, Zhao YH, Hu GF, Lv GJ. 2010. Effect of waterborne copper exposure on growth, hepatic enzymatic activities and histology in Synechogobius hasta. Ecotoxicol Environ Saf. 73(6):1286–1291.

Loro VL, Jorge MB, Rios K, Silva D, Wood CM. 2012. Oxidative stress parameters and antioxidant response to sublethal waterborne zinc in a euryhaline teleost Fundulus heteroclitus: Protective effects of salinity. Aquat Toxicol. 110(111):187–193.

Mackenzie NC, Brito M, Reyes AE, Allende ML. 2004. Cloning, expression pattern and essentiality of the high-affinity copper transporter 1 (ctr1) gene in zebrafish. Gene. 328(1–2):113–120.

Malekpouri P, Asghar A. 2011. Protective effect of zinc on related parameters to bone metabolism in common carp fish (Cyprinus carpio L.) intoxified with cadmium.: 187–196.

Matsuo AYO, Wood CM, Val AL. 2005. Effects of copper and cadmium on ion transport and gill metal binding in the Amazonian teleost tambaqui (Colossoma macropomum) in extremely soft water. Aquat Toxicol. 74:351–364.

Mcgeer JC, Szebedinszky C, Mcdonald DG, Wood CM. 2000. Effects of chronic sublethal exposure to waterborne Cu, Cd or Zn in rainbow trout. 1: Iono–regulatory disturbance and metabolic costs. Aquat Toxicol. 50:231–243.

Mebane CA, Dillon FS, Hennessy DP. 2012. Acute toxicity of cadmium, lead, zinc, and their mixtures to stream-resident fish and invertebrates. Environ Toxicol Chem. 31(6):1334–1348.

Mishra AK, Mohanty B. 2008. Acute toxicity impacts of hexavalent chromium on behavior and histopathology of gill, kidney and liver of the freshwater fish, Channa punctatus (Bloch). Environ Toxicol Pharmacol. 26(2):136–141.

Monteiro SM, Mancera JM, Fontaínhas-Fernandes A, Sousa M. 2005. Copper induced alterations of biochemical parameters in the gill and plasma of Oreochromis niloticus. Comp Biochem Physiol - C Toxicol Pharmacol. 141(4):375–383.

Monteiro SM, dos Santos NMS, Calejo M, Fontainhas-Fernandes A, Sousa M. 2009. Copper toxicity in gills of the teleost fish, Oreochromis niloticus: Effects in apoptosis induction and cell proliferation. Aquat Toxicol. 94(3):219–228.

Muhvich A, Jones R, Kane A. 1995. Effects of chronic copper exposure on the macrophage chemiluminescent response and gill histology in goldfish (Carassius auratus L.). Fish Shellfish….:251–264.

Naji T, Safaeian S, Rostami M, Sabrjou M. 2007. Toxic effect of zinc sulfate on gill tissues of common carp (Cyprinus carpio). J Environ Sci Technol. 9(2):29–36.

Nilsson GE, Dymowska A, Stecyk JAW. 2012. New insights into the plasticity of gill structure. Respir Physiol Neurobiol. 184(3):214–222.

Nimick DA, Harper DD, Farag AM, Cleasby TE, MacConnell E, Skaar D. 2007. Influence of in-Stream Diel Concentration Cycles of Dissolved Trace Metals on Acute Toxicity To One-Year-Old Cutthroat Trout (Oncorhynchus Clarki Lewisi). Environ Toxicol Chem. 26(12):2667.

Niyogi S, Nadella SR, Wood CM. 2015. Interactive effects of waterborne metals in binary mixtures on short-term gill-metal binding and ion uptake in rainbow trout (Oncorhynchus mykiss). Aquat Toxicol. 165:109–119.

Niyogi S, Wood CM. 2003. Effects of Chronic Waterborne and Dietary Metal Exposures on Gill Metal-Binding: Implications for the Biotic Ligand Model. Hum Ecol Risk Assess An Int J. 9(4):813–846.

Nunes B, Antunes • S C, Gomes • R, Campos • J C, Braga • M R, Ramos • A S, Correia • A T. 2015. Acute Effects of Tetracycline Exposure in the Freshwater Fish Gambusia holbrooki: Antioxidant Effects, Neurotoxicity and Histological Alterations. Arch Environ Contam Toxicol. 68:371–381.

Nunes B, Campos JC, Gomes R, Braga MR, Ramos AS, Antunes SC, Correia AT. 2015. Ecotoxicological effects of salicylic acid in the freshwater fish Salmo trutta fario: Antioxidant mechanisms and histological alterations. Environ Sci Pollut Res. 22(1):667–678.

Phuong LM, Huong DTT, Nyengaard JR, Bayley M. 2017. Gill remodelling and growth rate of striped catfish Pangasianodon hypophthalmus under impacts of hypoxia and temperature. Comp Biochem Physiol-Part A Mol Integr Physiol. 203:288–296.

Poleksic V, Mitrovic-Tutundzic V. 1994. Fish Gills as a Monitor of Sublethal and Chronic Effects of Pollution. p. 339–351.

Pratap HB, Wendelaar Bonga SE. 1993. Effect of ambient and dietary cadmium on pavement cells, chloride cells, and Na+/K+-ATPase activity in the gills of the freshwater teleost Oreochromis mossambicus at normal and high calcium levels in the ambient water. Aquat Toxicol. 26:133–149.

Qiu A, Hogstrand C. 2005. Functional expression of a low-affinity zinc uptake transporter (FrZIP2) from pufferfish (Takifugu rubripes) in MDCK cells. Biochem J. 390:777–786.

Ramesha AM, Guptha TRC, Katti RJ, Gowda G, Lingdhal C. 1996. Toxicity of cadmium to common carp Cyprinus carpio(Linn.). Environ Ecol. 14(2):329–333.

Rehwoldt R, Bida G, Nerrie B. 1971. Acute toxicity of copper, nickel and zinc ions to some hudson river fish species. Bull Environ Contam Toxicol. 6(5):445–448.

Reynders H, Van Campenhout K, Bervoets L, De Coen WM, Blust R. 2006. Dynamics of cadmium accumulation and effects in common carp (Cyprinus carpio) during simultaneous exposure to water and food (Tubifex tubifex). Environ Toxicol Chem. 25(6):1558–1567.

Ritz C, Streibig JC. 2005. Bioassay Analysis using R. J Stat Softw. 12(5).

Roberts RJ, Ellis AE. 2012. The pathophysiology and systematic pathology of teleosts. In: Roberts RJ, editor. Fish Pathology. Chichester: Wiley-Blackwell. p. 62–143.

Roopadevi H, Somashekar RK, Balagangadhar BR. 2011. Effects of pH on copper accumulation and toxicity in the common carp, Cyprinus carpio. J Environ Sci Eng. 53(3):335–340.

Santos DCM dos, Matta SLP da, Oliveira JA de, Santos JAD dos. 2012. Histological alterations in gills of Astyanax aff. bimaculatus caused by acute exposition to zinc. Exp Toxicol Pathol. 64(7–8):861–866.

De Schamphelaere KAC, Janssen CR. 2004. Bioavailability and Chronic Toxicity of Zinc to Juvenile Rainbow Trout (Oncorhynchus mykiss): Comparison with Other Fish Species and Development of a Biotic Ligand Model. Environ Sci Technol. 38:6201–6209.

Schwaiger J, Wanke R, Adam S, Pawert M, Honnen W, Triebskorn R. 1997. The use of histopathological indicators to evaluate contaminant-related stress in fish. J Aquat Ecosyst Stress Recover. 6(1):75–86.

Shahsavarani A, McNeill B, Galvez F, Wood CM, Goss GG, Hwang P-P, Perry SF. 2006. Characterization of branchial epithelial calcium channel (ECaC) in freshwater rainbow trout (Oncorhynchus mykiss). J Exp Biol. 209:1928–1943.

Shahsavarani A, Perry SF. 2006. Hormonal and environmental regulation of epithelial calcium channel in gill of rainbow trout (Oncorhynchus mykiss). Am J Physiol Regul Integr Comp Physiol. 291(5):R1490–8.

Spry DJ, Wood CM. 1985. Ion Flux Rates, Acid–Base Status, and Blood Gases in Rainbow Trout, Salmo gairdneri, Exposed to Toxic Zinc in Natural Soft Water. Can J Fish Aquat Sci. 42(8):1332–1341.

Spry DJ, Wood CM. 1985. Ion Flux Rates, Acid–Base Status, and Blood Gases in Rainbow Trout, Salmo gairdneri, Exposed to Toxic Zinc in Natural Soft Water. J Fish Aquat Sci. 42:1332–1341.

Spry DJ, Wood CM. 1989. A kinetic method for the measurement of zinc influx in vivo in the rainbow trout, and the effects of waterborne calcium on flux rates. J. Exp Biol. 142:425–446.

Stouthart XJHX, Haans JLM, Lock RAC, Bonga SEW. 1996. Effects of water pH on copper toxicity to early life stages of the common carp (Cyprinus carpio). Environ Toxicol Chem. 15(3):376–383.

Suiçmez M, Kayim M, Köseoglu D, Hasdemir E. 2006. Toxic effects of lead on the liver and gills of Oncorhynchus mykiss WALBAUM 1792. Bull Environ Contam Toxicol. 77(4):551–558.

Suresh A, Sivaramakrishna B, Radhakrishnaiah K. 1993. Patterns of Cadmium Accumulation in the Organs of Fry and Fingerlings of Freshwater Fish Cyprinus Carpio Following Cadmium exposure. Chemosphere. 26(5):945–953.

Suzuki N, Yamamoto M, Watanabe K, Kambegawa A, Hattori A. 2004. Both mercury and cadmium directly influence calcium homeostasis resulting from the suppression of scale bone cells: The scale is a good model for the evaluation of heavy metals in bone metabolism. J Bone Miner Metab. doi:10.1007/s00774-004-0505-3.

Thophon S, Kruatrachue M, Upatham ES, Pokethitiyook P, Sahaphong S, Jaritkhuan S. 2003. Histopathological alterations of white seabass, Lates calcarifer, in acute and subchronic cadmium exposure. Environ Pollut. 121(3):307–320.

Tishinova V. 1975. Study About the Toxic Action of Zinc on One Summer Old Carp: I. Lethal Concentrations,(In Bulgarian). God Sofii Univ Biol Fak 67 p 107–110, 1972/73(1975).

Triebskorn R, Telcean I, Casper H, Farkas A, Sandu C, Stan G, Colârescu O, Dori T, Köhler HR. 2008. Monitoring pollution in River Mureş, Romania, part II: Metal accumulation and histopathology in fish. Environ Monit Assess. 141(1–3):177–188.

Verma SR, Tonk IP, Dalela RC. 1981. Determination of the maximum acceptable toxicant concentration (MATC) and the safe concentration for certain aquatic pollutants. Acta Hydrochim Hydrobiol. 9(3):247–254. VMM. 2014. Geoview.

Vuorinen PJ, Keinänen M, Peuranen S, Tigerstedt C. 2003. Reproduction, blood and plasma parameters and gill histology of vendace (Coregonus albula L.) in long-term exposure to acidity and aluminum. Ecotoxicol Environ Saf. 54(3):255–276.

Wickham H. 2009. ggplot2: Elegant Graphics for Data Analysis. Springer-Verlag New York.

Wong CKC, Wong MH. 2000. Morphological and biochemical changes in the gills of Tilapia (Oreochromis mossambicus) to ambient cadmium exposure. Aquat Toxicol. 48:517–527.

Wood CM. 2001. Toxic responses of the gill. In: Shlenk D, Benson WH, editors. Target organ toxicity in marine and freshwater teleosts. London: Taylor and Francis. p. 1–101.

Wood CM, Farrel AP, Brauner CJ. 2012a. Homeostasis and Toxicology of Essential Metals. Farrel AP, Brauner CJ, editors.

Wood CM, Farrel AP, Brauner CJ. 2012b. Homeostasis and Toxicology of Non-Essential Metals.

Yancheva V, Velcheva –, Stoyanova –, Georgieva –. 2016. Histological Biomarkers in Fish As a Tool in Ecological Risk Assessment and Monitoring Programs: a Review. Appl Ecol Environ Res. 14(1):47–75.

